# Expanding Biomaterial Surface Topographical Design Space through Natural Surface Reproduction

**DOI:** 10.1101/2020.05.18.099663

**Authors:** Steven Vermeulen, Floris Honig, Aliaksei Vasilevich, Nadia Roumans, Aurélie Carlier, Manuel Romero, Paul Williams, Jorge Alfredo Uquillas, Jan de Boer

## Abstract

Surface topography guides cell behavior and is a tool to endow biomaterials with bioactive properties. The large number of possible designs makes it challenging to find the optimal surface structure to induce a specific cell response. The TopoChip platform is currently the largest collection of topographies with 2176 in silico designed micro-topographies. Still, it is exploring only a small part of the design space due to the boundary conditions of the design algorithm and the surface engineering strategy. Inspired by the diversity of natural surfaces, we assessed to what extend we could expand the topographical design space and consequently the resulting cellular responses using natural surfaces. To this end, we replicated twenty-six plant and insect surfaces in polystyrene and quantified their surface properties using white light interferometry, image analysis and principle component analysis. Next, we quantified mesenchymal stem cell morphology and the pattern of *Pseudomonas aeruginosa* colonization and compared it to previous data from TopoChip screens. Our data show that natural surfaces extended the TopoChip design space. Moreover, the natural surfaces induced MSC morphologies and bacterial attachment patterns not previously observed on the TopoChip. In the future, we will train our design algorithms with the results obtained by natural surface imprint experiments to further explore the design space and bio-active properties of surface topography.

## Introduction

Biological surfaces are interfaces between an organism and its environment and are the location where the organism deals with the chemical and physical reality of the outside world. Through evolution, bio-interfaces have acquired functional characteristics to support survival, which can be of a chemical nature, as seen in plant waxes that decrease moisture loss ^[1]^ and protect against UV radiation.^[2]^ On the other hand, topographical characteristics give surfaces very interesting and useful properties. For example, the setae on gecko feet allow movement on smooth vertical walls ^[3]^, and setae mimetics are used as dry and reversible adhesives for both robotic ^[4]^ and biomedical applications. ^[5]^ Superhydrophobicity, a material property observed on certain plant surfaces such as the Holy Lotus and Red Rose, is caused by hierarchical micro- and nanostructures, and results in self-cleaning surfaces.^[6,7]^ Mosquitos use specialized superhydrophobic nanostructures on their eyes to prevent the nucleation of fog droplets ^[8]^, and the tooth-like scales on shark skin provide drag reduction, anti-biofouling, and superoleophobicity which protects sharks against oil spills.^[9,10]^ Antimicrobial nano-structures on cicada wings may reduce the infection risk of implants ^[11,12]^ and through inspiration from the Nepenthes pitcher plant, lubrication fluids cover micro/nanostructured medical devices with repellent and self-cleaning surfaces.^[13–16]^ These fascinating evolutionary and bioengineered structures demonstrate the richness in natural surface topographical bioactivity and begs research to use natural surface topographical design to improve the performance of materials for industrial and clinical applications.

Surface topography is currently not only used as a design tool to improve biomaterials in prosthetic dentistry, tissue-engineering and regenerative medicine but also to enhance biocompatibility of medical devices.^[17–19]^ Surface topography has a very large design space, which we define as the universe of surface architectures, and ranges from randomly introduced roughness ^[20]^ to designed groove patterns.^[21]^ Additionally, topographies exist as pillars ^[22]^ or complex geometries ^[23,24]^, while curvature provides convex and concave shapes.^[25]^ All these structures, found both in micro- and nanometer dimensions, are known to affect the cells that are in contact with them.

The large topographical design space complicates the quest for the optimal topography for a specific application. To this end, high-throughput topography screening (HTS) platforms were developed and many novel bioactive surfaces have been discovered.^[26–29]^ However, HTS platforms still have their limitations because their design strategy only covers a small part of the whole design space. For example, structures in both nano- and micrometer dimensions are rarely present in the same platform and variable roughness levels are not included in a high-throughput setting. Man-made designed surface structures are frequently presented in an organized pattern, yet disorder also profoundly influences cell behavior.^[30,31]^ The bottleneck in producing a more diverse spectrum of surface topographies is not in the *in silico* design possibilities, where algorithms such as neural networks could aid the design and fine tuning of surface topographies^[24]^, but rather the technical limitations in surface topography manufacturing. Even though state of the art techniques such as two-photon stereo-lithography can handle the fabrication of complex shapes ^[32]^, its limited writing speed remains unsuitable for high-throughput applications. Photolithography, on the other hand, is mostly a 2D technique unsuitable to introduce design variability in height (z-axis) of complex topographies produced at large scale. We hypothesized that we can increase design space by using natural surfaces as a mold, and solvent casting as a microfabrication technique to replicate them into materials of interest, in our case tissue culture polystyrene. Many reports describe the replication of natural surfaces^[33–37]^ and in this work, we used a quantitative approach to compare design space coverage by artificial and natural surfaces.

We investigated the combination of multiple natural surfaces in one platform for the creation of a novel architectural design sub-space not commonly found in artificial platforms. We sampled a diverse set of 26 plant and animal surfaces, reproduced their structures in polystyrene, and investigated their potential for controlling cell behavior. We demonstrated that the structural diversity from the natural surfaces surpasses that of the micro-topographical TopoChip platform and show novel stem cell and bacterial bioactivity in this array of natural surface topographies.

## Results and Discussion

### Natural surface architectures exhibit a wide design variety

We selected sixteen plant and ten insect surfaces with a diverse set of surface properties based on reported phenomena such as super-hydrophobicity, anti-fouling or light reflection (see **Figure S1 and Figure S2**). Observation of the unprocessed specimens using scanning electron microscopy revealed an interesting variety in surface topography. For example, the Calla lily petal surface exhibited interconnected cuticular folds with ridges of 1 μm in height (**Fig. 1a, top row, left panel**), which is very different from the petal surface of the Red Rose, which has parallel-aligned hierarchical structures of 20 μm high micropapillae and nanofolds (**Figure 1a, top row, center panel**). Holy Lotus, known for its superhydrophobicity, has heptagonal inclinations, a nano rough surface and convex micro curvature resulting in a 10 μm high pillar in the center (**Figure 1a, top row, right panel**). On the rice surface, we find pillar structures with longitudinal ridges (**Figure 1a, bottom row, left panel)**. The *Huechys incarnata* wing is an interesting case, where merged pillars are present, which give rise to a variable structure size between 500 nm and 5 μm in diameter (**Figure 1a, bottom row, center panel**). Variable pillar sizes can be found on cicadas such as the *Yanga adriana*. (**Figure 1a, bottom row, right panel**). We classified plant surfaces in five groups: a) cuticular folds with low elevation, b) cuticular folds with high elevation, c) oriented structures, d) complex structures and e) micro-roughness (**Figure S3**). Three classes were observed on animal surfaces: a) nanopillars, b) pits, and c) curved surfaces (**Figure S4**).

**Figure 1.**
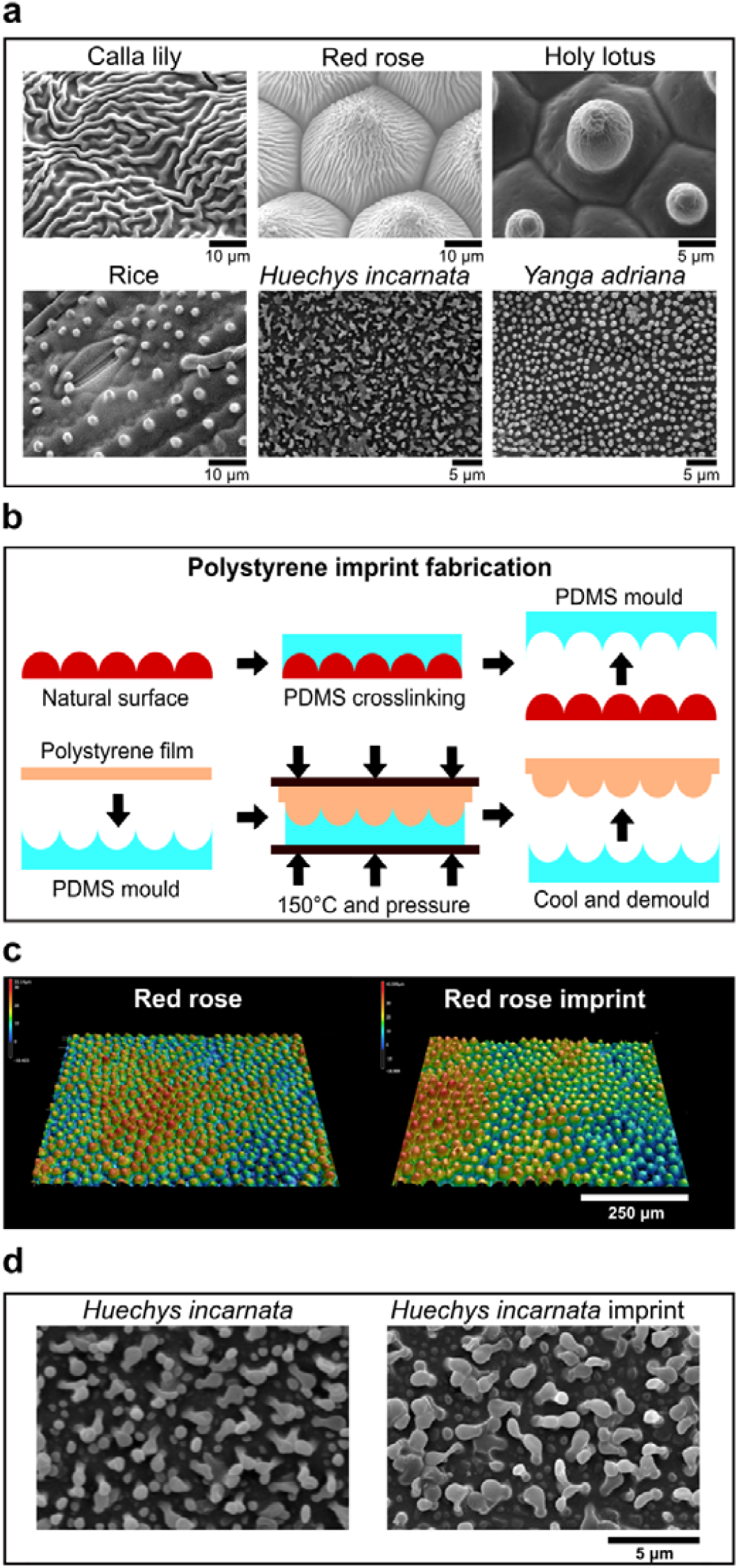
Natural surfaces exhibit large topographical variety. **a)** SEM images of leaves and insect wings utilized in this study. **b)** Fabrication scheme to imprint natural surface designs onto polystyrene. **c)** Leaf imprints can be transferred to polystyrene with high fidelity as seen in the profilometric images of Red Rose and its imprint counterpart. **d)** Insect wing imprints can be transferred to polystyrene with high fidelity as seen in the profilometric images of *Huechys incarnata* and its imprint counterpart.

### Natural surface architectures can be imprinted with high fidelity into polystyrene

We chose polystyrene as reference chemistry in which to compare cell-material interaction.^[38,39]^ In order to transfer the natural surface structures into polystyrene (PS), we used a relatively easy and fast technique that only requires the use of glass slides, binder clips, and a conventional oven (**Figure 1b**).^[40]^ Polydimethylsiloxane (PDMS) was cast upon the natural surface, after which it was cured at room temperature for 24 h. The PDMS containing the negative imprint of the natural topographies was peeled off the natural surface (**Figure 1b, top row** and **Figure S5A**) and a ‘sandwich’ was created containing glass slides, Teflon sheets, the PDMS imprint and a polystyrene sheet (**Figure 1b, bottom row** and **Figure S5B**). The construct was placed in the oven and after one hour, the construct was cooled to room temperature and the PS imprint peeled off the PDMS.

We next compared the profilometric images of the PS imprints and original specimens and found using high-resolution microscopy that the imprints were successfully transferred with high fidelity from natural surfaces into PS. Examples are provided for the leaves of the Red Rose (**Figure 1c)**, and *Huechys incarnata* (**Figure 1d**). Sub-micron structures present on the cuticular folds of the Red Rose and Holy Lotus were also successfully replicated (**Figures S6 and S7**).

After observing the correct imprinting of natural surface topographies onto PS, we assessed as proof of concept the superhydrophobic properties of both the Red Rose and Holy Lotus imprints by measuring the water contact angle. No significant differences were seen between the natural and PS imprints (**Figure S8**). The rose petal effect, resulting in pinning of a water droplet, was demonstrated by inverting and tilting the imprint. The Lotus effect, which is characteristic of the rapid rolling of water droplets on the surface was also seen on the PS Holy Lotus imprint (data not shown). These examples demonstrate that also the biophysical properties of natural topographies can be replicated on PS.

### Natural surface architectures occupy a different part of the design space and induce distinct cell morphology compared to TopoChip micro-topographies

The objective of this study was to determine whether natural surface topographies occupy a different part of the design space than our previously established library of randomly generated TopoChip topographies.^[23,41]^ We quantified the dimensional features of 199 randomly sampled TopoChip surfaces and the twenty-six natural surfaces based on the design file and interferometry data, respectively and plotted the differences using principal component analysis (PCA). (**Figure 2a**).^[23]^ PCA is a dimension reduction technique that condenses multi-dimensional data into fewer features and allows a visual representation of the variation of the principal components. Each data point in the PCA plot represents a single surface and the distance between dots represent their dissimilarity. Interestingly, TopoChip topographies formed two distinct and separate clusters: one mainly distinguished by large features and smooth, round edges (**Figure 2b**) and another mainly with small features and rectangular edges (**Figure 2d)**. On the other hand, natural surfaces formed a third ‘linear’ cluster (**Figure 2c**) with micro- and nanotopographies, demonstrating that natural surfaces represent an unexplored part of the design space.

**Figure 2.**
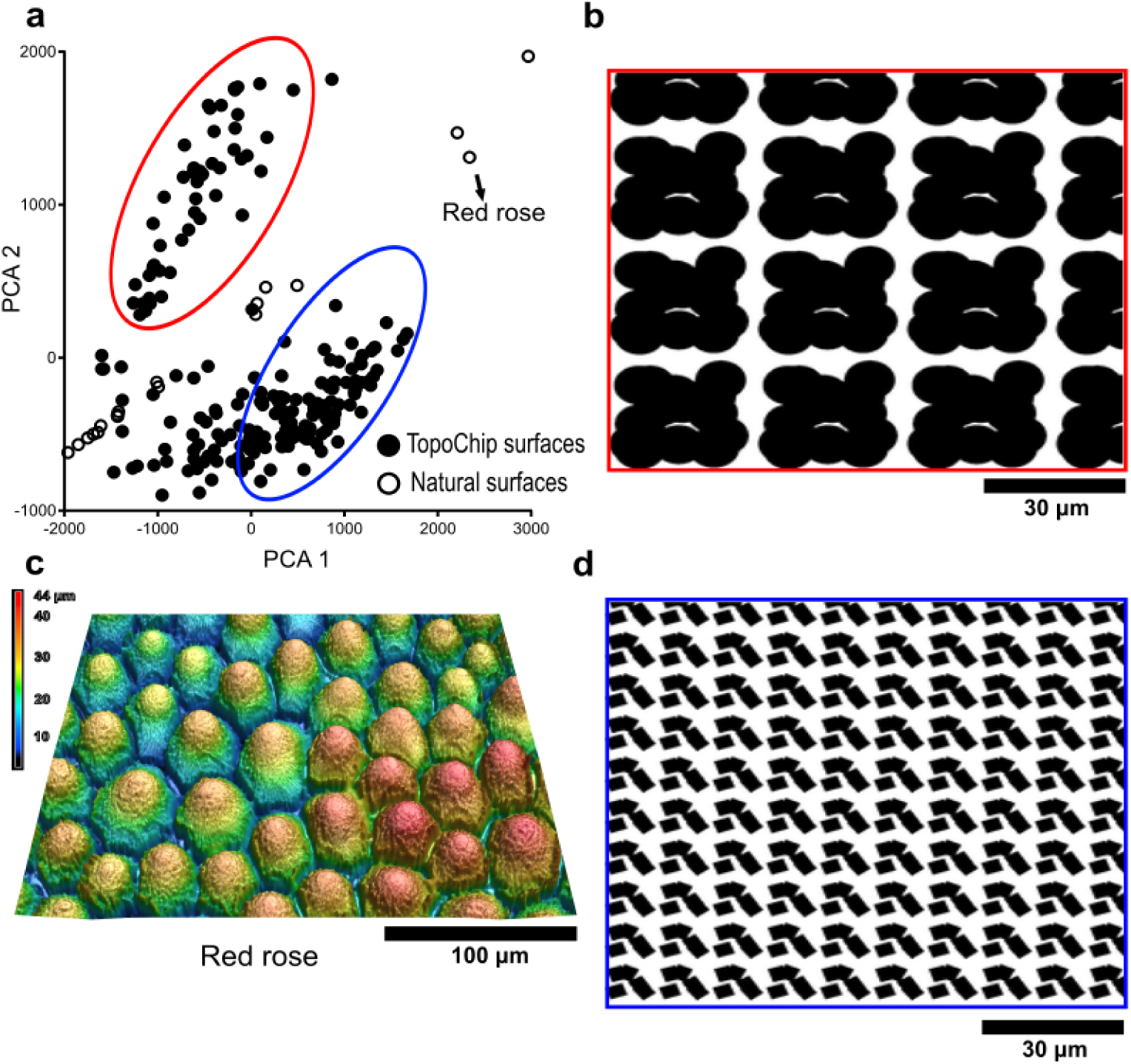
Natural surfaces expand the topographical design space. **a)** Principal component analysis of the TopoChip and natural surfaces. Note three distinct clusters: the red cluster with its representative CellProfiler topography in **b)**, and the blue cluster with its representative CellProfiler topography in **d)**. The third ‘linear’ cluster is represented by the profilometric topography of Red Rose (**c**).

Cell shape and surface topography are highly correlated^[42–44]^, and we wondered if a larger design space also leads to yet unobserved cell shape features. To this end, we seeded human mesenchymal stem cells (hMSCs) onto the PS imprints of 26 natural surfaces and onto 28 TopoChip topographies which were selected to capture the whole range of cell shape variation on the TopoChip.^[45]^ F-actin and DNA were stained, and quantitative information of cell and nucleus area, and cell compactness and solidity data were extracted for all surfaces (**Figure 3**).^[46]^ In **Figure 3a**, the effect of topography on MSC cell and nucleus area is compared relative to MSCs on flat PS. The further a data point is from the unit coordinates and closer to the plot origin, the smaller is the cell and nucleus area. In general, hMSCs seeded on natural surfaces show substantially larger cell and nucleus areas compared to the size of cells on the TopoChip (**Figure 3a**), and these cell and nucleus sizes fall between those corresponding to flat and TopoChip surfaces. Representative fluorescence images of flat, onion, and TopoChip surface 11 facilitate the visualization of size differences (**Figure 3c)**.

**Figure 3.**
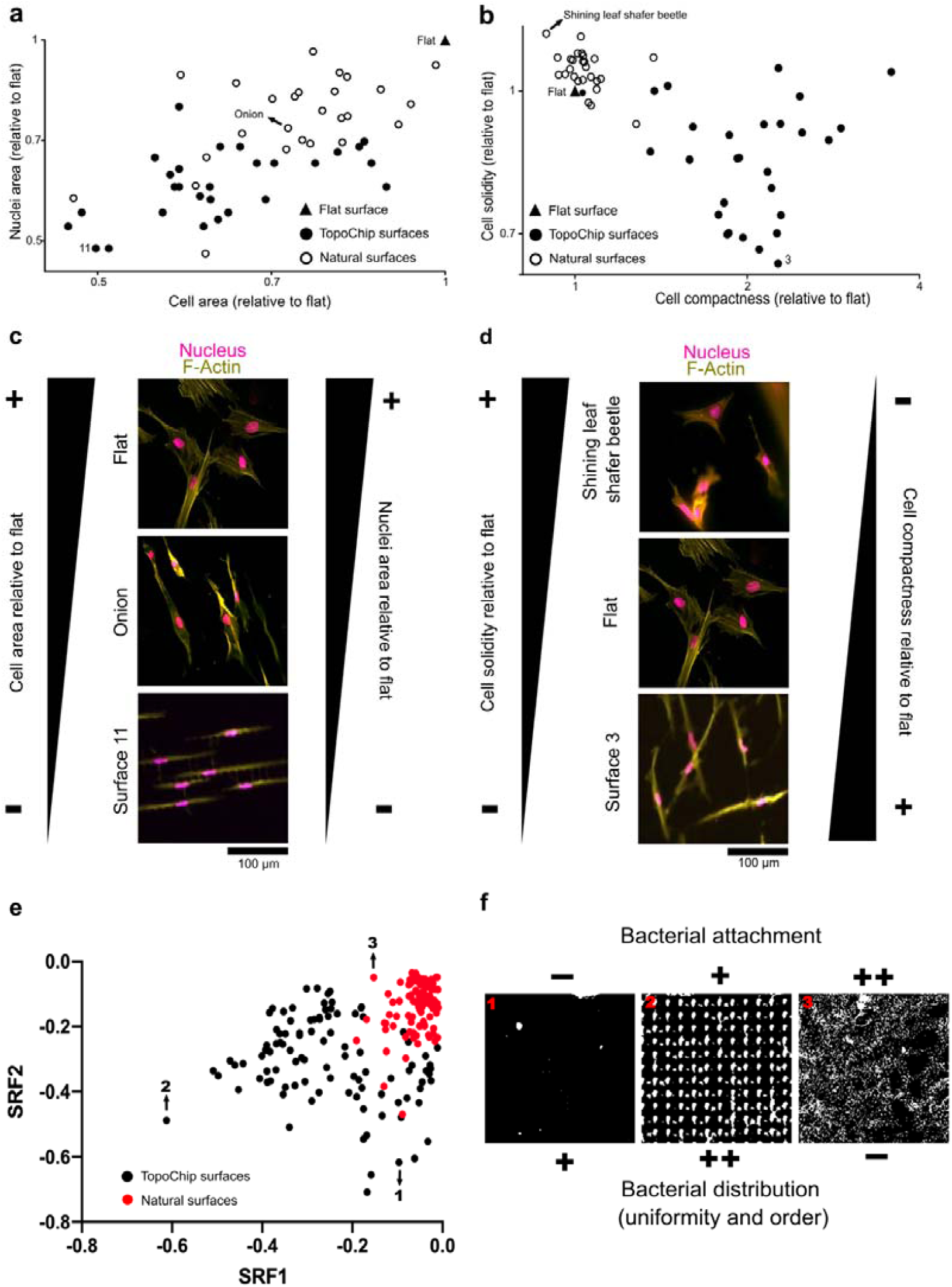
TopoChip and natural surfaces have a quantitative different effect on the shape of mesenchymal stem cells and the attachment distribution of *P. aeruginosa*. **a)** Cell and nucleus area quantification of hMSCs on TopoChip and natural surfaces. **b)** Cell compactness and solidity quantification of hMSCs on TopoChip and natural surfaces. **c)** Visual representation of the change in cell and nucleus area of hMSCs on TopoChip and natural surfaces. **d)** Visual representation of the change in cell compactness and solidity of hMSCs on TopoChip and natural surfaces. **e)** Spatial relationship feature plot of *P. aeruginosa* on TopoChip (points 1 and 2) and natural surfaces (point 3). **f)** Visual representation of the bacterial attachment and distribution of *P. aeruginosa* on TopoChip (image 1 and 2) and natural surfaces (image 3). (-) = low, (+) medium, (++) high. A random selection of natural surfaces was replicated in polystyrene and introduced in the TopoChip to expand the topographical design space to ultimately influence cell behavior. Natural surfaces show features in the micro- and nanoscale, that are not found in the artificially designed TopoChip, that uniquely influence size, shape, and attachment distribution of human mesenchymal stem cells and *Pseudomonas aeruginosa*.

Cell compactness relates to cell elongation, and cell solidity is inversely correlated to cell branching and radial filopodia protrusion. A high compactness value indicates a more elongated cell, and a high solidity value indicates that the cell is less branched and has less filopodial protrusions extending radially (**Figure 3b and Figure 3d**). We observed that cells on natural surfaces tend to have less cell branching and radial filopodia protrusion, with most cells on natural surfaces with solidity values of more than 1. TopoChip surfaces typically induce compactness values of more than 1, as seen on the TopoChip surface 3, where cells were more elongated and branched than cells on natural surfaces (**Figure 3d**). In general, the analysis of cell compactness and solidity between designed and natural surfaces shows that natural topographies induce cell morphologies distinct from those observed on the TopoChip.

### Natural surface topographies induce distinct profiles of Pseudomonas aeruginosa colonization

Bacteria colonize diverse natural and artificial surfaces forming biofilms that confer protection against environmental stresses.^[47]^ Such biofouling is highly problematic in both industrial and medical contexts. Bacterial biofilm formation can be prevented by incorporating toxic biocidal agents, by modifying surface chemistry or surface topography.^[48,49]^ Several studies have shown that bacterial attachment can be controlled using patterned surfaces featuring repeating topographical elements of sizes ranging from nanometers to micrometers. Some of these patterned surfaces have been based on non-fouling natural surfaces including shark skin, plant leaves and insect wings.^[48,50]^ However, to our knowledge no systematic high throughput screens have yet been published. Hence, we compared the patterns of bacterial attachment to the 26 natural surfaces with designed (TopoChip) imprints. *P. aeruginosa* was chosen as the model bacterium as it is an environmentally ubiquitous micro-organism that readily adheres to natural and engineered surfaces and is highly problematic in a clinical infection context as it forms multi-antibiotic resistant biofilms on implanted medical devices.^[51]^ The attachment distribution profiles of Pseudomonas aeruginosa after 4 h growth on the natural surfaces were compared with those observed on 100 randomly sampled PS TopoChip topographies using the texture parameters in CellProfiler software. The spatial relation features (SRF) 1 and 2 are shown in **Figure 3e**, which clearly demonstrated that the natural surfaces occupy a different area of distribution to the TopoChip surfaces. In general, we noted more uniform and ordered bacterial distribution on the TopoChip surfaces (**Figure 3f**).

Finally, we are designing experiments to extract individual features of artificial and natural surfaces to induce a specific cell response and use this chip as a screening platform for studying patient-specific innate inflammatory response to implants and coating materials.

## Conclusion

In this work, we transferred features of natural surfaces to cell culture platforms to increase the topographical design space. We demonstrated that natural surfaces (i) can be transferred with high fidelity in polystyrene, (ii) occupy an unexplored area of topographical design space relative to the TopoChip, (iii) uniquely alter the size and shape of human mesenchymal stem cells, and (iv) affect the spatial distribution of *Pseudomonas aeruginosa* attachment. The TopoChip design strategy centers around the design parameters that feed the TopoChip design algorithm and as such, will only give TopoChip hits, which in the context of the topographical landscape can be seen as a local minimum. With natural surfaces we can step out of our local minimum and bring new and unique unit features into the design algorithm. Our current algorithm does not allow such modifications which is why we are exploring the use of neural networks as a novel topography design tool. In this manuscript we used only twenty-six natural surfaces, a minuscule subset of all the plants and insects on the planet. We envision the replication of the topography of plants and insects in the most biodiverse regions on Earth. ^[52]^ For instance, we would like to explore topographical design space at the Tiputini Biodiversity Station in the Ecuadorian Amazon rainforest, one of the most biodiverse tropical forests in the world^[53–56]^, and scan natural surfaces with a portable profilometer based on interferometry to obtain digital representation of natural surfaces. ^[57]^ Deep learning algorithms developed in house^[58,59]^ and elsewhere^[60]^ can decouple complex topographical information of natural surfaces and provide guides in selecting surfaces that cover uncharted topographical territory. Two-photon lithography can then be used to generate the surfaces and analyze their bio-active properties. Our work is ongoing to unveil advanced and nature-inspired surfaces with unique bio-instructive properties that could be used in disease and patient specific regenerative medicine applications.

## Acknowledgements

SV, NH, NR, JAU and JdB acknowledge the financial support of the Dutch province of Limburg. SV is grateful for the support of the European Union’s Horizon 2020 Programme (H2020-MSCA-ITN-2015; Grant agreement 676338). This work was also supported by the Engineering and Physical Sciences Research Council, UK (grant no. EP/N0016615/1) and the Biotechnology and Biological Sciences Research Council, UK (grant no. BB/R012415/1). AC kindly acknowledges the Dutch province of Limburg in the LINK (FCL67723) (“Limburg Investeert in haar Kenniseconomie”) knowledge economy project and a VENI grant (number 15075) from the Dutch Science Foundation (NWO). The authors thank the botanical garden “Hortus Botanicus” of Leiden, the Netherlands, for the donation of plant species, and Mr. Louay Waked for his help in material fabrication. SV kindly acknowledges Daniel Pereira for providing advice in fabrication and material production.

Received: ((will be filled in by the editorial staff))

Revised: ((will be filled in by the editorial staff))

Published online: ((will be filled in by the editorial staff))

## Extending Surface Topographical Design Space through Natural Surface Reproduction

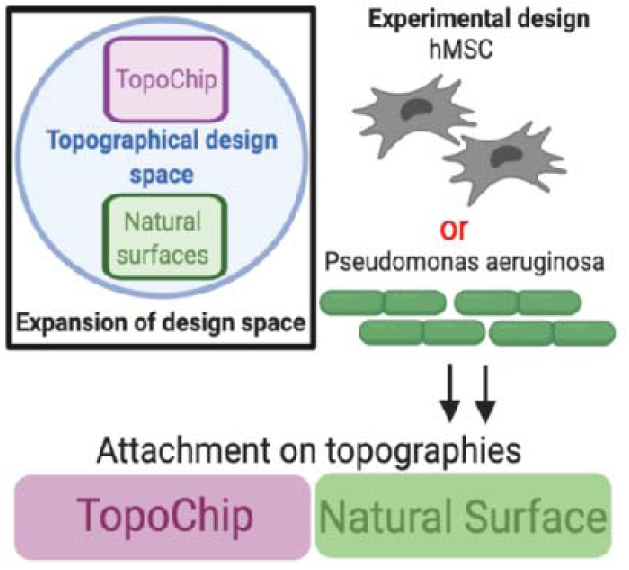

## Supporting Information

### Experimental Section

#### Fabrication of artificial micro-topographies in the TopoChip

A detailed description describing the fabrication of the micro-topographical surfaces can be found elsewhere. ^[61]^ In brief, the inverse pattern was etched on a silicon wafer by directional reactive ion etching (DRIE). To facilitate subsequent demolding procedures, the silicon waver was coated with a layer of perfluorooctyltrichlorosilane (FOTS, Sigma-Aldrich). Polydimethylsiloxane (PDMS; Sylgard® 184 Elastomer Kit, Dow Corning) in a 10:1 silicon/curing agent ratio (w/w) was cured on the waver at 85 °C for 6 h to generate a positive mold. OrmoStamp hybrid polymer (micro-resist technology Gmbh) was subsequently poured on the PDMS imprint together with a borofloat waver containing a layer of Ormoprime (micro-resist technology Gmbh). UV treatment allowed the cross-linking of a final negative Ormostamp imprint, which is used as a template for the hot embossing step at 140 °C for 10 min to generate a polystyrene imprint.

#### Fabrication of natural surface topographies in the TopoChip

Dried insects used in this study were purchased from the supplier “The Bugmaniac” (http://www.thebugmaniac.com). The Holy Lotus (Nelumbo nucifera), Rice (Oryza sativa), Plaintain Lily (Hosta sp.), and Hardy Canna (Thalia dealbata) were kindly donated from the botanical garden “Hortus Botanicus” of Leiden (https://www.hortusleiden.nl). Other plants were purchased at the local flower-shop. Replication of natural surfaces was achieved through a two-step fabrication process. ^[40]^ The first step included the fabrication of a polydimethylsiloxane (PDMS) mold, which contains the negative imprint of the natural surface. A sample of the fresh natural surface of choice was fixed onto a petri dish using double-sided adhesive tape (Double Fix, Bison). The silicon resin and curing agent (Sylgard® 184 Elastomer Kit, Dow Corning) were mixed in a 10:1 ratio (w/w) and cast onto the natural surface. Subsequently, the liquid mixture was degassed and cured for 48 h at room temperature to prevent heat damage. Finally, the solid PDMS mold was separated from the natural surface. In the second step, the PDMS mold served as a template for transferring the structures into PS using hot embossing. To achieve this, an assembly was made in which a 190 μm thick PS film was placed on the PDMS mold clamped between two Teflon sheets and microscope slides to apply constant pressure. To cross the glass-transition of the PS film, the entire system was inserted in an oven for 1h at 150 °C. Afterward, the assembly was removed from the oven and cooled down to room temperature to allow the PS film to solidify. Finally, the PS surface with transferred surface topography was carefully peeled from the PDMS mold.

#### Scanning electron microscopy imaging

Scanning electron microscopy was used for examination of the surface topography of natural surfaces and quality control of their respective PS replicas. Preparation of fresh natural samples consisted of fixation (2% glutaraldehyde in 0.1M cacodylate buffer) for 1hr. Subsequently, the samples were washed three times in cacodylate (0.1M) before dehydration by immersion in a graduated series of ethanol in water (50%, 70%, 90%, 100%). Then, samples were critical point dried (EM CPD300, Leica) using 15 exchange cycles with slow gas out and heating speed settings. All samples were mounted on SEM stubs using carbon conductive adhesive tape. Finally, samples were sputter-coated with gold (Sputter Coater 108auto, Cressington) for 100 seconds prior to imaging using a scanning electron microscope (XL-30, Philips) at 10kV.

#### Cell culture

Adipose-derived human mesenchymal stem cells (AD-hMSCs) were purchased from Lonza, which were isolated from a 42-year-old female. Basic medium for AD-hMSCs consisted of MEM Alpha GlutaMAX (Gibco), fetal bovine serum (FBS, 10% v/v), ascorbic-acid-2-phosphate (ASAP, 0.2 mM), and penicillin/streptomycin (10 U mL^-1^). Cells were grown at 37 °C in a humid atmosphere at 5% CO_2_. For experimental purposes, cells were seeded on flat and topographical surfaces at a density of 10000 cells/cm^2^ unless stated otherwise.

#### Fluorescent imaging

After cell culture, cells were washed with phosphate-buffered saline (PBS, Sigma-Aldrich) before fixation in formaldehyde (3.6% v/v) for 10 min at 37°C. After, cells were washed three times with PBS and permeabilized by the addition of Triton-X-100 (0.1% v/v) in PBS for 10 min. Following this, samples were blocked using serum (1:100) in PBT (0.02% Triton-X-100, PBS and 0.6% bovine serum albumin) for 1hr. After another washing step, samples were incubated for 1 h in Phalloidin with a fluorochrome attached to visualize F-actin (1:500, ThermoFisher). Finally, Hoechst 33258 (1:1000, ThermoFisher) was used for visualization of nuclei. After three subsequent washing steps, the samples were mounted in Mowiol.

#### Human mesenchymal stem cell morphology analysis

Fixed and stained samples were inverted, and fluorescent images were acquired through the glass coverslip using a fully automated Nikon Eclipse Ti-U microscope in combination with an Andor Zyla5.5 four megapixels camera. Fluorescent images were analyzed through CellProfiler 3.1.8 ^[46]^, applying custom-made pipelines. All images were cropped in order to remove out-of-focus objects. Objects touching the border of the subsequent images were filtered out of the dataset. After illumination corrections, the morphology of the nucleus was captured by the Otsu adaptive thresholding method applied on the Hoechst 33258 image channel. Subsequently, cell morphology was determined by applying propagation and Otsu adaptive thresholding on the Phalloidin image channel. Mis segmentation artifacts were removed by applying an arbitrary threshold on nuclei and cell size. For visualization purposes, we enhanced the brightness and contrast of the images representing cellular morphologies. The imaging software Fiji was used for image visualization.^[62]^

#### Bacteria imaging and data acquisition

*Pseudomonas aeruginosa* PAO1 was chosen to test performance of natural topography replicas against bacterial surface attachment. PAO1 was routinely grown at 37°C in lysogeny broth (LB) or LB agar. Tryptic soy broth (TSB) supplemented with human serum (10% v/v) was used as the growth medium for bacterial attachment assays. Prior to incubation with bacteria, topographies were washed by dipping in distilled water and sterilized in ethanol (70% v/v). The air-dried samples were placed in petri dishes (60 mm x 13 mm) and incubated statically at 37°C in 10 ml of growth medium inoculated with diluted (optical density: OD600 nm = 0.01) bacteria from overnight cultures. After 4h incubation, topography samples were removed and washed in PBS (pH 7.4) to remove loosely attached bacterial cells. After rinsing with distilled water, attached cells were stained with SYTO9 (50 µM; Molecular Probes, Life Technologies) for 30 min at room temperature. After staining, topographies were rinsed with distilled water, air-dried and mounted on a glass slide using Prolong antifade reagent (Life Technologies). Topographies were then imaged by confocal laser scanning using a Zeiss LSM 700 microscope (Carl Zeiss, Germany) and 488 nm laser as light source. Since the bacterial cells may attach at different heights on the micro-patterns, images were initially acquired as ∼75 µm range Z-stacks (1.5 µm steps - 50 slides) from the TUs using a 10x objective (Zeiss, EC Plan-Neofluar 10x/0.30 Ph 1).

#### PCA analysis

Greyscale images of the natural surfaces were captured through profilometric imaging through a Keyence VK-H1XM-131 profilometer. Height profiles were represented as grayscale images, where black represented the bottom of the surfaces and white top. For artificial surfaces, pixel values 0 correspondent to the bottom and 10 to the top of the pillars, which matches their height in µm. Since the natural surfaces height profile gradually varied from the bottom to the top of the surface, the pixel values also were counted and were arranged in a range from 0 (bottom) to the height value as measured by profilometry. As the input for the algorithm, we have used pixel values directly, 250000 values per surface. Resulting images contained information about the height of the elements as well as their structure on the area of 200×200 µm with resolution 500×500 pixels. Profilometry software was further used to correct for sample tiltness. We further corrected images by applying local contrast enhancement algorithm as implemented in sci-kit Python 3.7.3 package. Intensity values in the image were normalized further by subtracting minimum pixel value and dividing by max pixel intensity value per surface. The final image was obtained by multiplying a resulting matrix by height value as measured with the profilometer.

#### Texture quantification

Spatial distribution of bacteria attachment on the TopoChip and natural surfaces was quantified by image texture features, which were extracted from the thresholded images. Image analysis was performed in CellProfiler 3.1.8.

#### Texture Feature Selection

To distinguish texture features that were the most discriminative between TopoChip and natural surfaces, we employed a binary classification algorithm. Particularly we trained the Extreme Gradient Boosting (EGB) model on a hold-out subset that contained over 500 images of the bacteria attachment on either TopoChip or natural surfaces. The accuracy of the trained model was validated using the training set and the accuracy value was 0.98. An accuracy of 1.0 corresponds to a perfectly performing classification algorithm. We further quantified the importance of the features with the Mean Decrease Accuracy (MDA) algorithm that measures how model accuracy decreases when a feature is excluded from the prediction.

#### Spatial relationship features

We identified that the texture feature ‘InfoMeas1 for scales 64 and 2’ were the most important for distinguishing bacteria attachment between TopoChip and natural surfaces. ‘InfoMeas1’ is a measure of the total amount of information contained within a region of pixels derived from the recurring spatial relationship between specific intensity values.^[63]^. We rename them in the text and Figure 3e as Spatial Relationship Feature 1 (SRF1) and Spatial Relationship Feature 2 (SRF2). For visualization purposes, we transformed SRF1 by taking its cubic root. The scatter plot in Figure 3f represents 100 randomly selected surfaces from both TopoChip and natural groups.

**Supplementary Figure 1.**
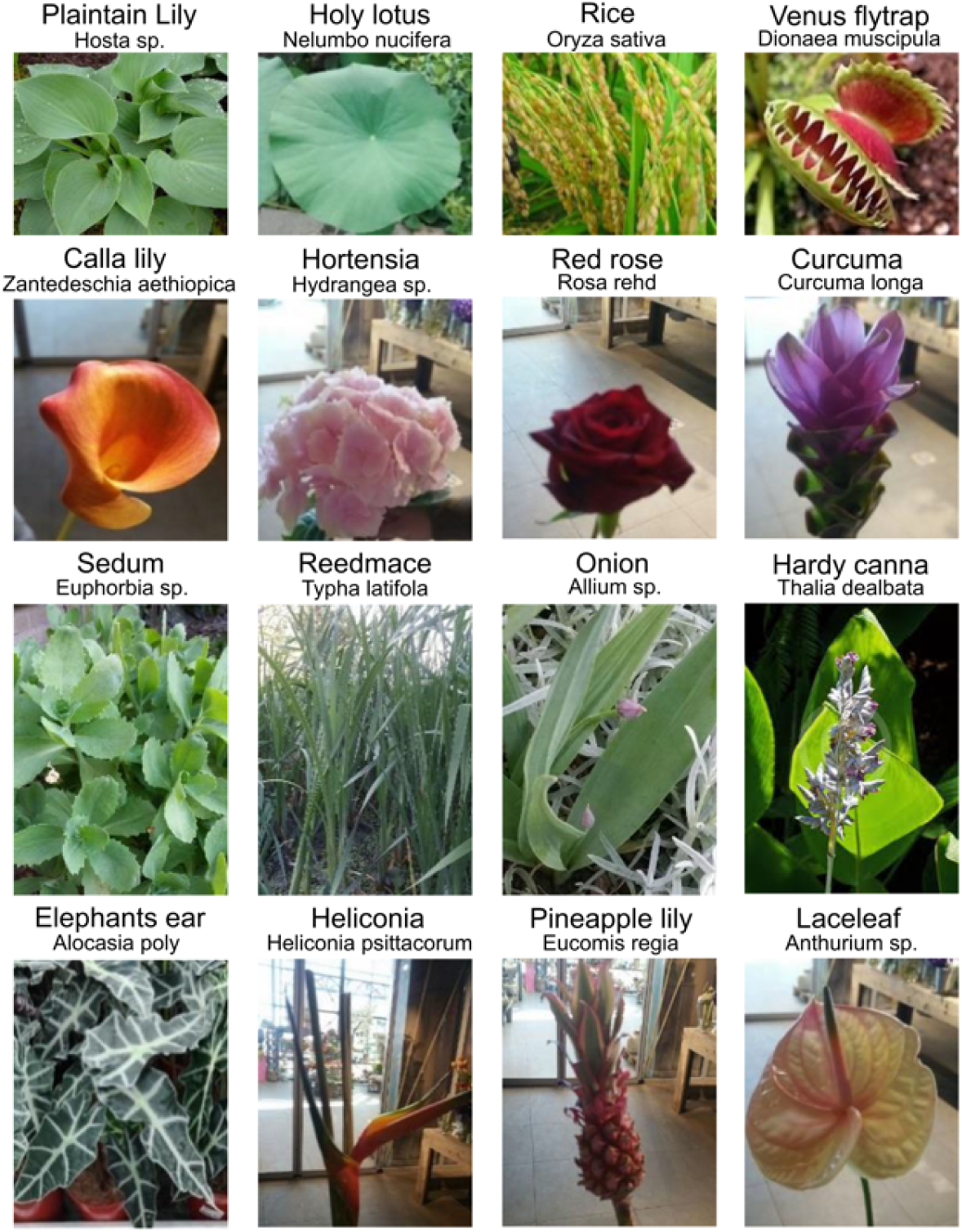
Photographs of the sixteen plants used to obtain PS replicas of their leaf’s topography.

**Supplementary Figure 2.**
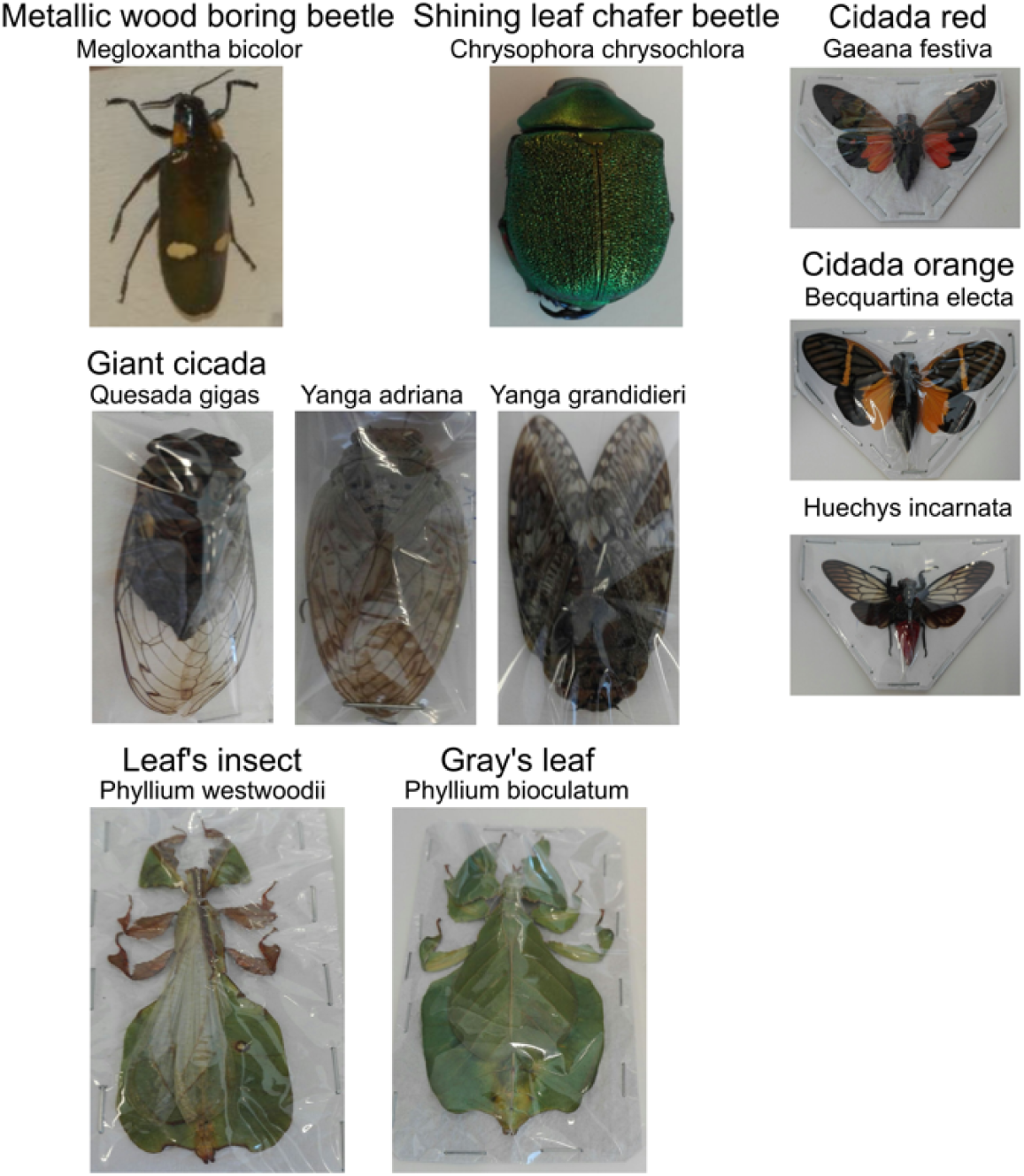
Photographs of the ten insects used to obtain PS replicas of their wings.

**Supplementary Figure 3.**
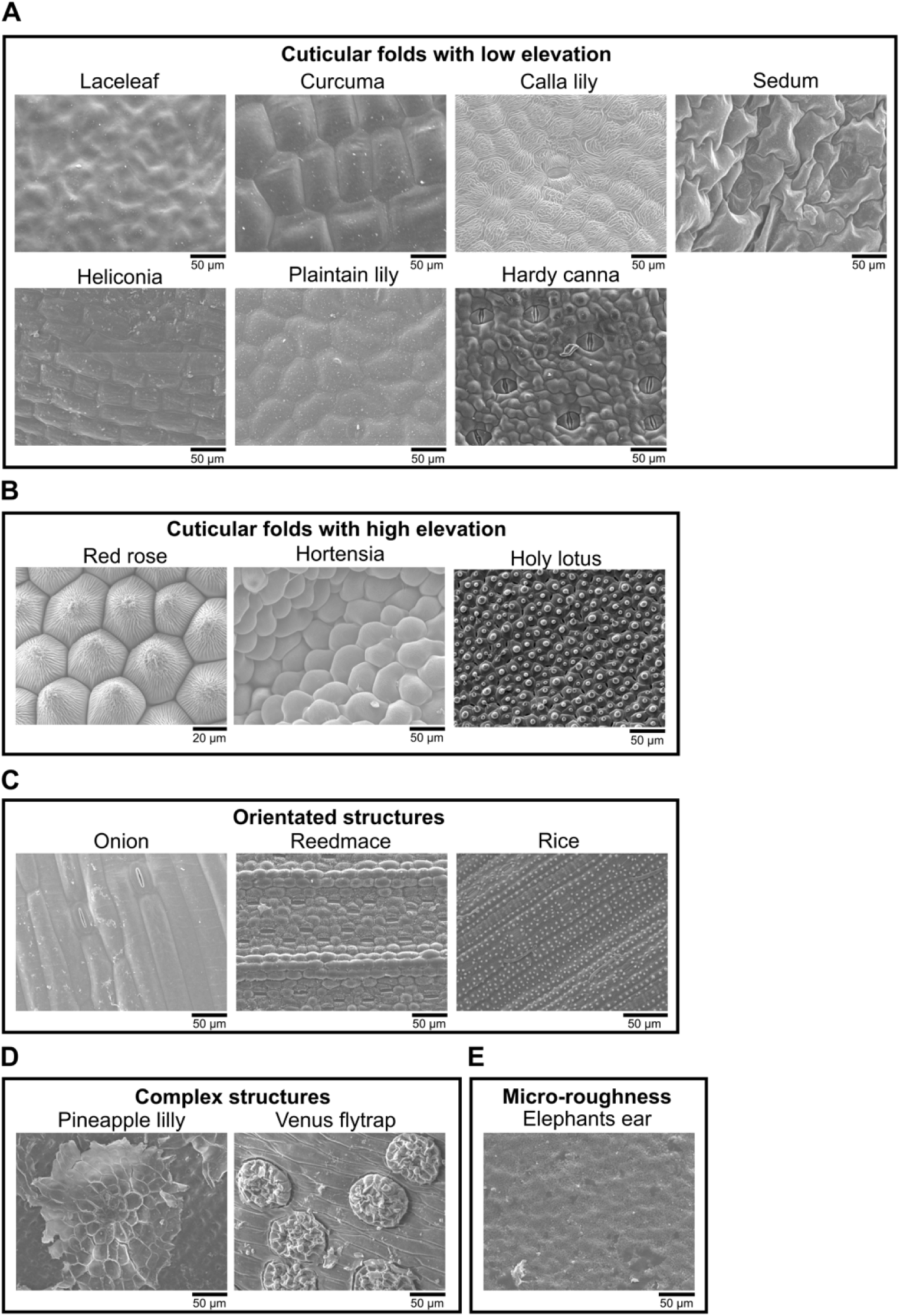
SEM images of plant surfaces revealed a broad diversity in surface topography. **A)** Structures with low elevation are present on the Laceleaf, Calla lily, Curcuma, Sedum, Heliconia, Plantain lily, and Hardy canna. **B)** Cuticular folds with complex and hierarchical structures are found on the Red rose, Hortensia, and Holy lotus. **C)** Oriented structures are found on the Onion, Reedmace, and Rice. **D)** Complex structures are found on the Pineapple lilly and Venus flytrap. **E)** Crystalline waxes present on the Elephants Ear give rise to micro-roughness structures.

**Supplementary Figure 4.**
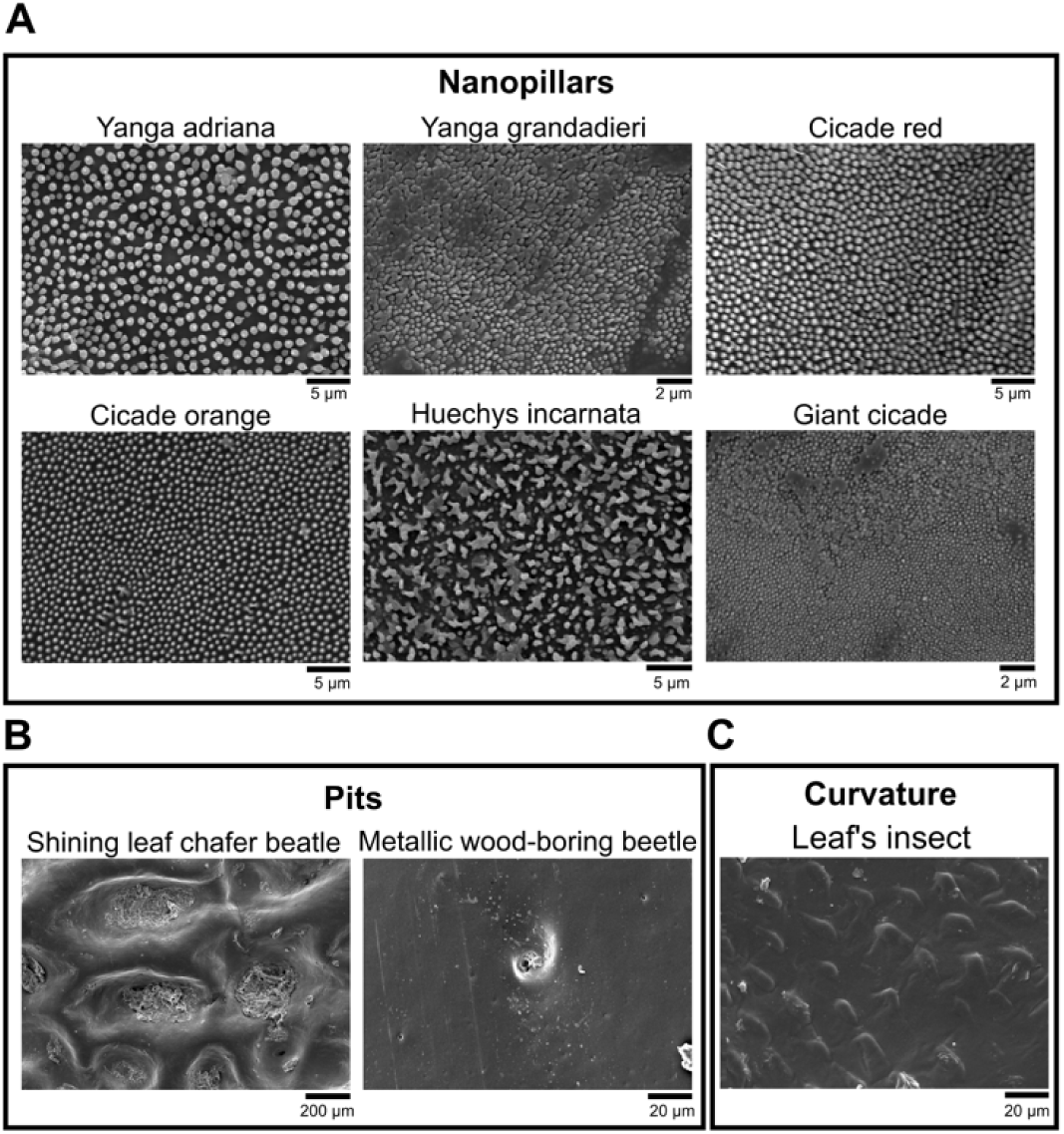
SEM images of insect surfaces expanded the design space with nanotopographies, wells, and curvature. **A)** Nanopillars with different dimensions are found on different cicada species. **B)** Beetles exhibit pit-like structures on their surface, as shown for the Shining leaf chafer beetle and the metallic-wood boring beetle. **C)** Surfaces containing curvature are found on insects resembling leaves of plants, as shown for the Leaf’s insect.

**Supplementary Figure 5.**
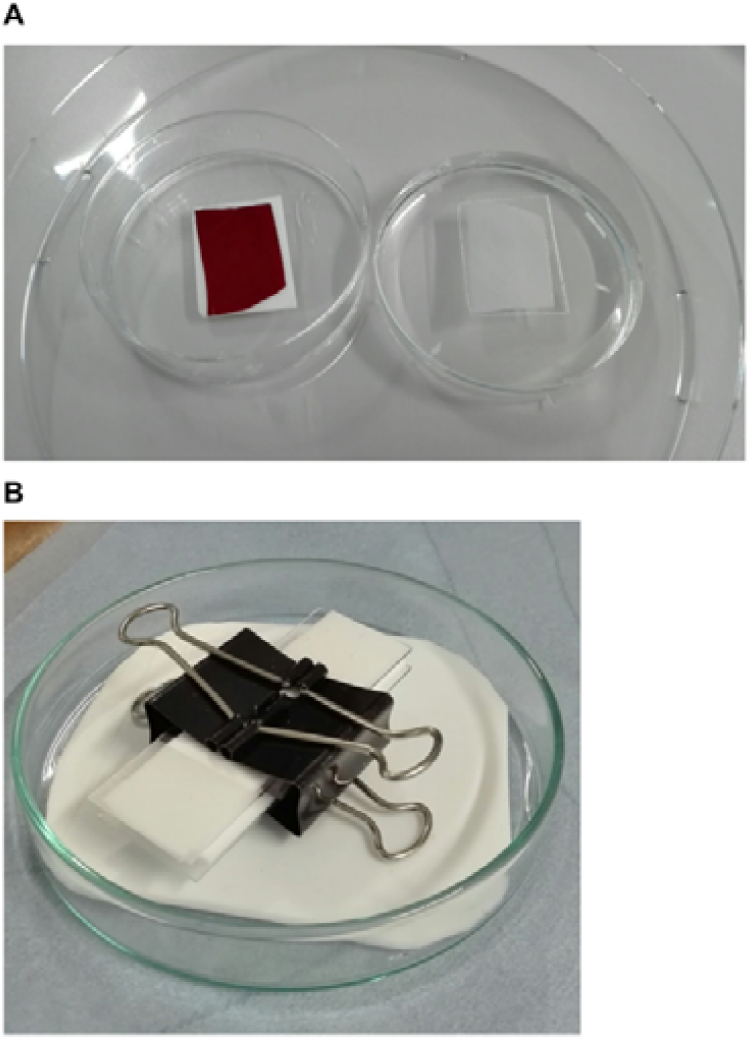
Easy ‘sandwich’ imprinting fabrication. **A)** PDMS is poured on the Red rose imprint, after which it is peeled-off after curing for 24h. **B)** Afterwards, Teflon sheets, PDMS, and PS are sandwiched and pressured together with binders before placing the ‘sandwich’ in the oven.

**Supplementary Figure 6:**
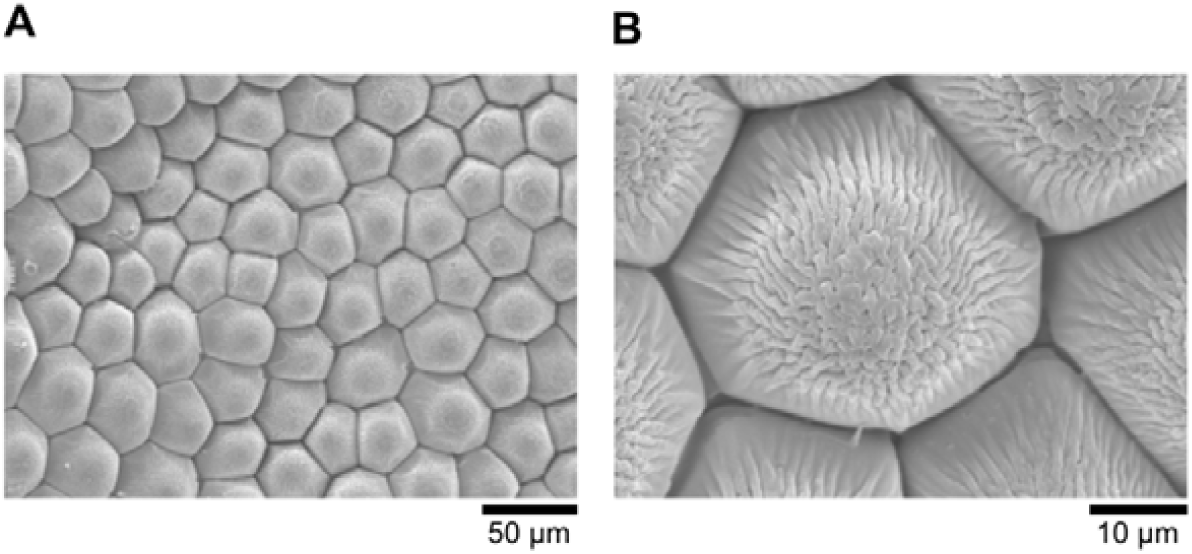
The Red Rose topography can be transferred with high fidelity to polystyrene. SEM images of the Red Rose polystyrene imprint at **A)** low, and **B)** high magnification.

**Supplementary Figure 7:**
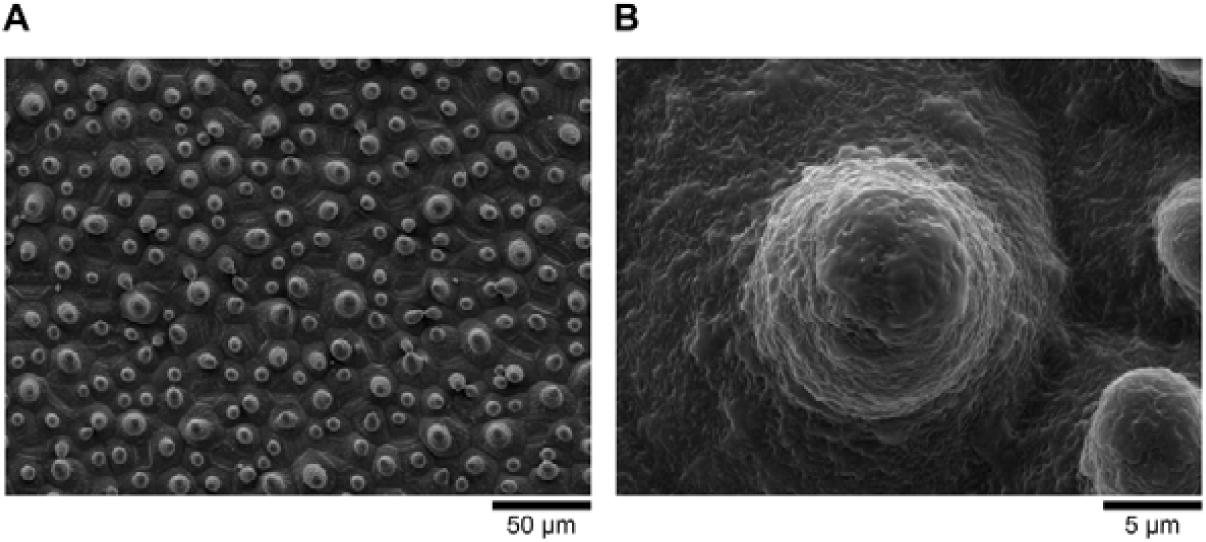
The Holy Lotus topography can be transferred with high fidelity to polystyrene. SEM images of Holy Lotus polystyrene imprint at **A)** low, and **B)** high magnification.

**Supplementary Figure 8:**
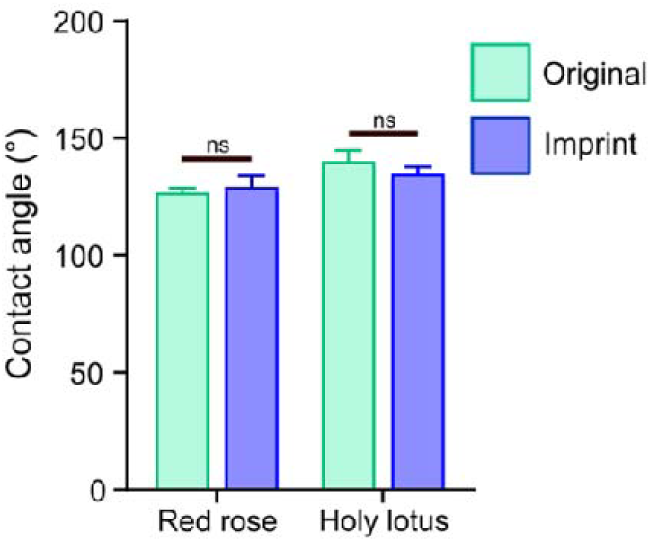
No difference in the water contact angle was seen between the original surface and the PS imprints of Red Rose and Holy Lotus.

